# Leveraging Pretrained Deep Protein Language Model to Predict Peptide Collision Cross Section

**DOI:** 10.1101/2024.09.11.612388

**Authors:** Ayano Nakai-Kasai, Kosuke Ogata, Yasushi Ishihama, Toshiyuki Tanaka

**Affiliations:** Graduate School of Engineering, Nagoya Institute of Technology, Nagoya, Aichi 466-8555, Japan; Department of Molecular Systems Bioanalysis, Graduate School of Pharmaceutical Sciences, Kyoto University, Kyoto 606-8501, Japan; Laboratory of Proteomics for Drug Discovery, National Institute of Biomedical Innovation, Health and Nutrition, Ibaraki, Osaka 567-0085, Japan; Graduate School of Informatics, Kyoto University, Kyoto 606-8501, Japan

**Author notes:** **Corresponding authors:** Correspondence to Yasushi Ishhama or Toshiyuki Tanaka.

## Abstract

Collision cross section (CCS) of peptide ions provides an important separation dimension in liquid chromatography/tandem mass spectrometry-based proteomics that incorporates ion mobility spectrometry (IMS), and its accurate prediction is the basis for advanced proteomics workflows. This paper describes novel experimental data and a novel prediction model for challenging CCS prediction tasks including longer peptides that tend to have higher charge states. The proposed model is based on a pretrained deep protein language model. While the conventional prediction model requires training from scratch, the proposed model enables training with less amount of time owing to the use of the pretrained model as a feature extractor. Results of experiments with the novel experimental data show that the proposed model succeeds in drastically reducing the training time while maintaining the same or even better prediction performance compared with the conventional method. Our approach presents the possibility of prediction in a “greener” manner of various peptide properties in proteomic liquid chromatography/tandem mass spectrometry experiments.

## 1 Introduction

Proteins are important biological elements responsible for various functions of living organisms, and a systematic understanding of when, where, and how these proteins are expressed is necessary for systemwide analysis of biological functions [1]. Therefore, an important issue in proteomics is how to efficiently identify and quantify the vast number of proteins present in cells and tissues [2]. Recent advances in liquid chromatography/tandem mass spectrometry (LC/MS/MS) have significantly improved the coverage of bottom-up proteomics [3]. However, a typical human proteome sample consists of more than 10 million protease-digested peptides [4], whose complexity is beyond the separation capabilities of current LC/MS/MS systems [5].

Recently, ion mobility spectrometry (IMS) has gained attention as yet another promising separation method to be combined with LC/MS/MS [6, 7, 8, 9, 10]. IMS separates molecules in terms of their charge and shape by measuring the mobility of ions moving in a buffer gas flow under the influence of an electric field [11]. The frequency of ion-gas collisions, also known as the collision cross section (CCS), determines the ion mobility in the gas phase [12]. Thus, even ion species of the same *m/z* may exhibit different CCS values due to different conformations they take [13]. The extended separation space provided by ion mobility resolves various problems caused by the insufficient separation of peptide ions in the conventional LC/MS/MS. For example, it can improve the separation of peptide isomers, which are peptides with the same sequence but different positions of post-translational modifications. Additionally, the improvement of peptide separation can lead to better quantitation accuracy [7, 9, 14]. IMS is not only effective for improving the separation efficiency of peptide ions, but it also has the potential for improving peptide identification [15, 16]. While peptide identification primarily relies on MS/MS spectra, utilizing additional information such as peptide retention time can aid in the identification process, particularly for data from target acquisitions or data-independent acquisitions. However, accurate prediction of peptide retention time is necessary for approaches utilizing retention time information to be effective [17, 18]. It is also the case with those utilizing IMS data: For better analysis of proteomic IMS data, it would be desirable if one can accurately predict CCS values of peptide ions. Several groups have so far proposed CCS prediction algorithms. Clemmer and coworkers established the model called intrinsic size parameter (ISP), which represents the relative size of each amino acid residue in a peptide sequence [13, 19, 20]. While this model works for a specific set of peptides, it has inherent limitations due to the use of the peptide’s amino acid compositions but not the sequences. This ISP parameter has been further expanded to incorporate some sequence information [16].

On the other hand, inspired by great successes of deep learning in various research fields, the use of a deep neural network (NN) model for the CCS prediction problem has been proposed in [21], where a bidirectional LSTM model [22], trained from scratch with a dataset of 559, 979 unique peptide ion data, was used for CCS prediction. However, these models are extensively trained with the peptide sequence of less than 30 amino acid residues, and mostly with the doubly charged species, which are typically observed in proteomic experiments. The prediction of CCS values of longer peptides is challenging due to the limited availability of data and their enhanced structural variability. Longer peptides tend to have higher charge states, such as triple, quadruple charges and more, which results in higher variability in CCS among species even within similar *m/z* ranges. Therefore, accurate prediction of CCS for longer peptides requires both new experimental data and new prediction models.

Our proposal for CCS prediction is to use a pretrained deep protein language model as a feature extractor from peptide sequences, and train a separate NN, which we call a prediction NN, to predict CCS values on the basis of the extracted features. The overall model architecture of our proposal, which we name the pretrained protein language model-based network (PPLN), is depicted in Fig. 1. It has been argued [23] that a deep (natural) language model trained with a large corpus of a language implicitly acquires grammatical information of that language. Likewise, a pretrained deep protein language model trained with a large-scale database of protein sequences is believed to acquire structural information of proteins [24] (so called “protein grammar”). Indeed, it has been demonstrated in [25] that the feature representation provided by a pretrained deep protein language model named evolutionary scale modeling1b (ESM-1b) is useful in prediction of secondary structure and residue-residue contacts in proteins. As the CCS value of a peptide ion is regarded as being determined by the conformation of the ion particles in the drift tube, one can expect that the features obtained by such a deep protein language model that encodes structural information of proteins will be useful in prediction of CCS values of peptide ions as well.

**Figure 1:**
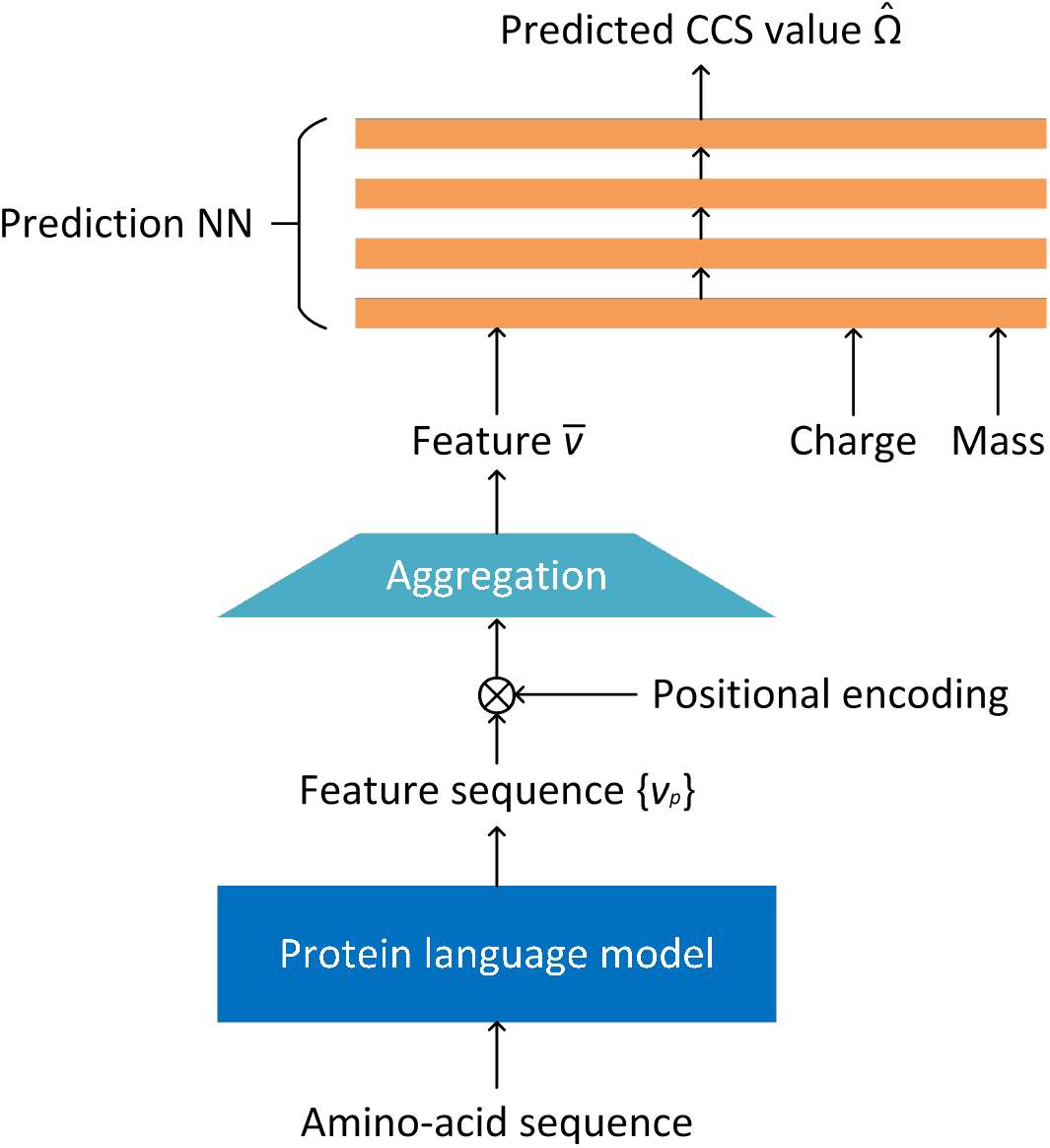
Model architecture of PPLN for prediction of CCS value. An amino-acid sequence is input into a pretrained protein language model. Positional encoding is applied to the obtained feature sequence. The feature sequence is aggregated to form a fixed-size feature, which is then input into the prediction NN along with charge and mass.

Use of a pretrained protein language model as a feature extractor for CCS prediction might limit performance of CCS prediction compared with the approach of training a dedicated complex deep NN model from scratch, when the quality of the extracted features by the pretrained model is not good. It will have several advantages, on the other hand, when the quality of the features is good. It will make the CCS prediction problem easier to solve: One can use a simpler prediction NN, and train it with a smaller-sized training dataset and with less amount of time. Training of the prediction NN will thus be performed in a “greener” manner than training a dedicated complex NN model from scratch, on cheaper computer hardware and with less energy consumption.

In this study, we used a newly measured CCS dataset containing longer peptides to investigate how effective PPLN is in improving accuracy compared with traditional deep learning models and how “green” the steps required to create a CCS predictor can be performed.

## 2 Results

### 2.1 Model architecture of PPLN

A number of deep protein language models for general purpose [25, 26, 27, 28, 29, 30, 31] and specific tasks [32, 33] have so far been proposed. Such a deep protein language model, especially one for general purpose, can be used as a feature extractor, by removing the output layer of the model and regarding the outputs of the pre-output layer as a feature representation of the input. Although we used ESM-1b as the feature extractor of PPLN in our experiments, any pretrained protein language model can in principle be used as the feature extractor. A feature extractor that is based on a deep protein language model typically takes as its input a variable-length amino-acid sequence, and outputs a sequence of features, whose length is the same as the length of the input sequence. One then has to aggregate the variable-length feature sequence into a fixed-size representation to feed it into the prediction NN, as shown in Fig. 1.

Although aggregation with simple averaging as used in the original ESM-1b would work well in certain tasks as in [25], we found that introduction of aggregation considering amino acid positions, i.e., positional encoding (PE) in the aggregation, worked better than simple averaging. The fixed-size feature representation, along with the charge number and the mass of the ion, is then fed into the prediction NN, which outputs a prediction 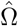 of the CCS value Ω of the input peptide ion. We included the charge number and the mass as the input of the prediction NN because the masses of peptides obviously have a strong correlation with the CCS values, and the charge numbers of peptide ions also have a strong influence on the CCS values. Inclusion of the charge number and the mass to the input of the prediction NN is thus expected to facilitate the learning of the prediction NN.

We next discuss our design of PE. The CCS value of a peptide ion can be affected by several factors. Among them, one can expect that amino acid residues located near the terminals have stronger effects than those located in the central part of the peptide, as suggested, e.g., by the length-specific multiple linear regression (LS-MLR) study [16]. We thus designed our PE in the aggregation stage in such a way that it reflects features of those residues located near either of the terminals of a peptide ion, rather than the absolute positions of the residues relative to the N-terminal, the latter of which we call the unidirectional PE in this paper.

More concretely, assume that we use a feature extractor which, when fed with a length-*l* peptide ion sequence, outputs a length-*l* sequence 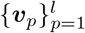 of *d*-dimensional feature vectors. We then propose the following PE, which we term the bidirectional PE: The feature vector sequence 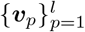 is aggregated into a single 2*d*-dimensional vector 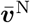 via

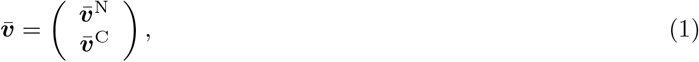

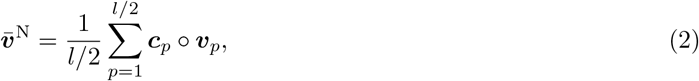

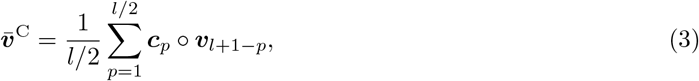

where ° denotes the element-wise (Hadamard) product of vectors, and where ***c***_*p*_ = (*c*_*p*,1_, …, *c*_*p,d*_)^⊤^ is the encoding vector for position *p* from one of the terminals. That is, 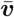 summarizes the features from the half of the amino acid sequence on the N-terminal side, and 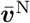 summarizes the features from the other half of the amino acid sequence on the C-terminal side. (When *l* is odd, one may have to introduce an appropriate rounding of the half-integer *l/*2. In our implementation used in the experiments, we used the python built-in function `round`().) These vectors are concatenated to form the resulting feature vector 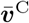 One can argue that the bidirectional PE bears the same spirit as that in [21] where they used the bidirectional LSTM rather than the (unidirectional) LSTM to deal with peptide sequences. As mentioned above, an alternative choice to the bidirectional PE might be the unidirectional PE, where the *d*-dimensional aggregated vector 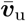 is obtained via

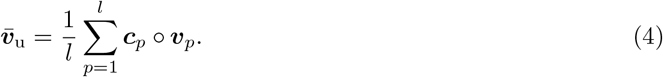

We will experimentally compare performance of the model with the bidirectional PE and that with the unidirectional PE in the later section.

### 2.2 Dataset construction

The emergence of proteomic IMS technologies and the availability of large-scale peptide CCS data have significantly improved the performance of peptide CCS prediction [15, 16]. However, the current proteomic data mainly cover peptides with a length of less than 30 amino acids, which limits our understanding of the behaviors of longer peptides that are of general interest. To address this gap in knowledge and to improve CCS prediction for longer peptides, we constructed an experimental peptide CCS dataset using phosphoproteome data. Phosphopeptides are known to have more missed cleavages and tend to be longer than non-phosphopeptides [34]. To obtain a unique set of peptides, we digested HeLa cell extracts with seven proteases (trypsin, LysargiNase, Lys-C, Lys-N, Glu-C, Asp-N, and chymotrypsin) as described previously [16], enriched phosphopeptides from the digests [35], and fractionated the resulting phosphopeptides with SCX-StageTip [36]. We then dephosphorylated the phosphopeptides using calf intestine alkaline phosphatase [34] to generate non-phosphorylated peptides with more missed cleavages. Mascot automated database search algorithm was used to identify the peptides, and IonQuant [37] was used to extract the peptide precursor ion features (mass, retention time, ion mobility, and intensity). We filtered out peptide ions bearing variable modifications and only considered the most abundant feature for each peptide ion. Phosphoproteomes contain distinct, longer sequences compared with global proteome samples (Fig. S1).

The dataset consists of 91,677 unique peptide ion data. It includes 11 % of singly charged, 57 % of doubly charged, 25 % of triply charged, and 7 % of quadruply charged ions, which are shown in Fig. 2(a). Frequencies of peptide C-terminal and N-terminal amino acids are also summarized in Fig. 2(b) and Fig. 2(c), respectively. Figure 3 shows a plot of the experimental CCS values Ω versus the *m/z* values. The experimental CCS values and *m/z* range from 289 to 1162 Å^2^ and 381 to 1798, respectively.

**Figure 2:**
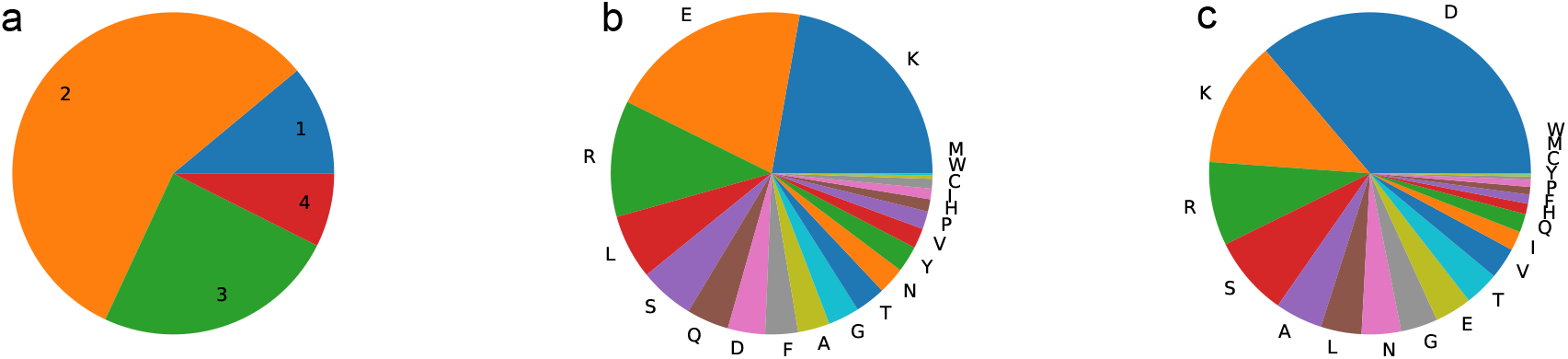
Summary statistics for the peptide dataset prepared from HeLa lysate using seven proteases. **a** Frequency of peptide charge numbers. **b** Frequency of peptide C-terminal amino acids. **c** Frequency of peptide N-terminal amino acids.

**Figure 3:**
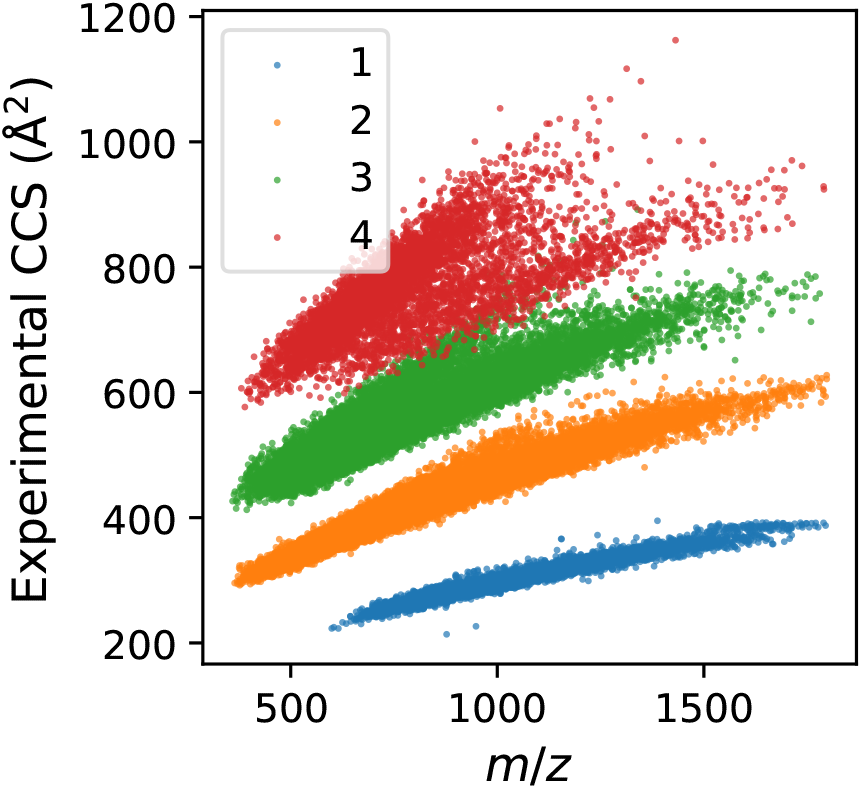
Distribution of 91,677 unique peptide ion. Experimental CCS values of singly, doubly, triply, and quadruply charged ions versus *m/z*.

### 2.3 Experimental model settings based on pretrained deep protein language model

This section introduces detailed settings of the prediction model PPLN in the experiments.

We used the pretrained ESM-1b model [25] as the feature extractor in PPLN, whose output is a sequence of *d* = 1280-dimensional feature vectors. As for the encoding vectors ***c***_*p*_ used in PE, we considered the following functional form

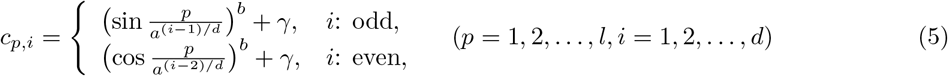

with parameters (*a, b, γ*). This formulation is inspired by the positional encoding used in the attention mechanism of the transformer architecture for a deep natural language model [38]. We used (*a, b, γ*) = (1000, 1, 0) in the following experiments.

The dataset mentioned in the previous section was randomly divided in two parts, where 73,342 ions (80 % of the total) for training and the remaining 18,335 (20 %) for testing. We used PyTorch and adopted minibatch learning with batch size 200. The number of layers of the prediction NN was set to 10 and their dimensions were 1000 except for the last layer that outputs the scalar CCS value. We trained the prediction NN with Adam optimizer (learning rate: 0.0003) and the MSE loss function using the training data over 400 epochs, and tested the CCS prediction performance of the trained model on the test data.

### 2.4 Evaluation of prediction error from and correlation with experimental CCS values

Figure 4 shows scatter plots of the predicted CCS values obtained by the proposed PPLN. Figure 4(a) includes results of predicted CCS values versus experimental CCS values and two statistics, root mean squared error (RMSE) and Pearson’s correlation coefficient *r*. The definitions are given by

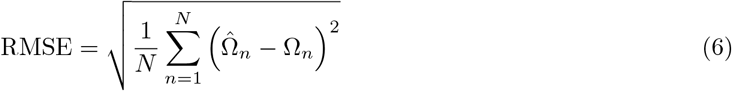

and

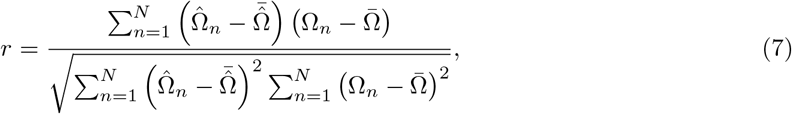

respectively, where *N* = 18, 335 is the number of test samples, where Ω_*n*_ and 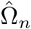 are the experimental and predicted CCS values, respectively, of *n*th test sample, and where 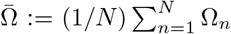 and 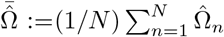 are the averages of the experimental and predicted CCS values. This figure also includes the results of LS-MLR [16] and the bidirectional LSTM-based method [21]. The deep NN used in the bidirectional LSTM-based method was trained from scratch using the training dataset of this paper, rather than the pretrained model using the dataset provided in [21]. The predicted CCS values by LSMLR tended to overestimate the experimental CCS values, especially for ions with larger CCS values. It is worth noting that the distribution of larger CCS peptides was split into two populations: One with overestimated CCS values predicted by LS-MLR and the other with fairly predicted CCS values. This can be mostly explained by the length of the peptides: The peptides with overestimated CCS were longer than the other ones (Fig. S2). On the other hand, the proposed PPLN provided better predictive performance with the lower RMSE and the higher correlation coefficient. Figures 4(b) and 4(c) show the relation to *m/z* and length of the ions. These results indicate that the CCS prediction of longer peptide ions with higher *m/z* is difficult. For the establishment of a better CCS prediction model, improving the prediction accuracy of these ions is mandatory.

**Figure 4:**
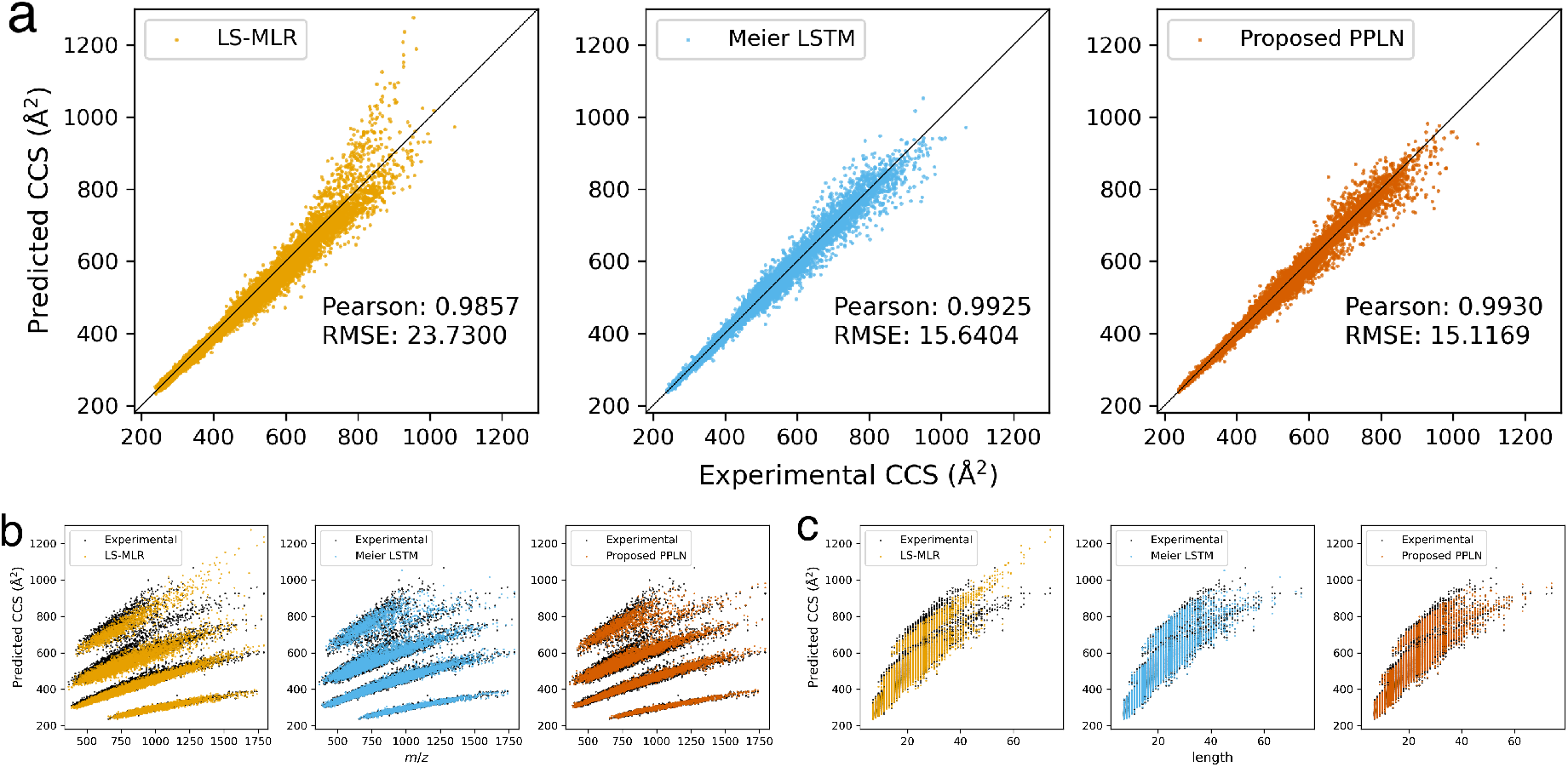
Scatter diagrams and statistics in terms of predicted CCS values. **a** Predicted CCS values vs. experimental CCS values including Pearson’s correlation coefficient and RMSE. **b** Predicted CCS values vs. *m/z*. **c** Predicted CCS values vs. length.

The predicted CCS values and the two statistics are summarized by charge number in Fig. 5. From the figure, we can see that the predictions for triply and quadruply charged ions were more difficult than the singly and doubly charged ions because the performance of all the three methods compared became worse as the charge number was higher. We can also see in Fig. 5(d) that the peptides with CCS values overestimated by LS-MLR are typically quadruply charged. This is consistent with the fact that longer peptides tend to have higher charge states. The proposed PPLN achieved the best performance among the methods in all the cases. The results imply the high applicability of the proposed method.

**Figure 5:**
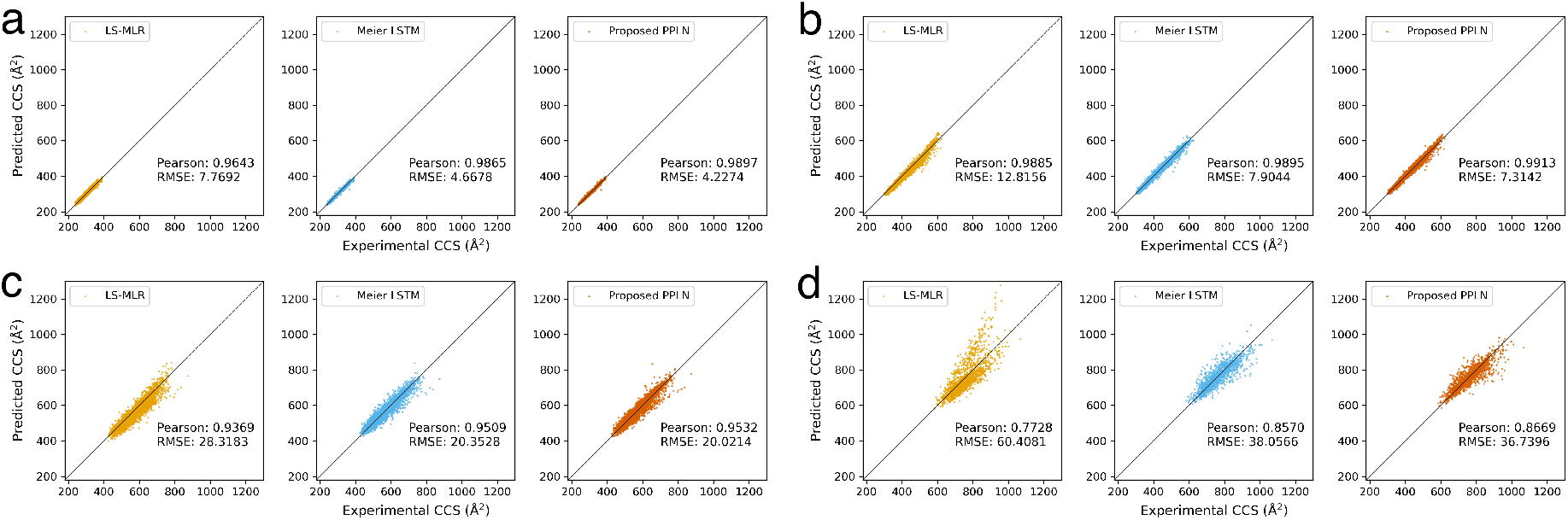
Predicted CCS values and statistics by charge. **a** Scatter diagram for singly charged ions including Pearson’s correlation coefficient and RMSE. **b** For doubly charged ions. **c** For triply charged ions. **d** For quadruply charged ions.

### 2.5 Ablation study

In this section, we verify necessity of the components of the proposed PPLN via ablation study, where we compared the predictive performance of the proposed method with the same method except that a part of the components was removed from it, in order to see if the removed part was important. Table 1 summarizes RMSE and Pearson’s correlation coefficient *r* obtained by the proposed method and methods with the removal.

**Table 1:**
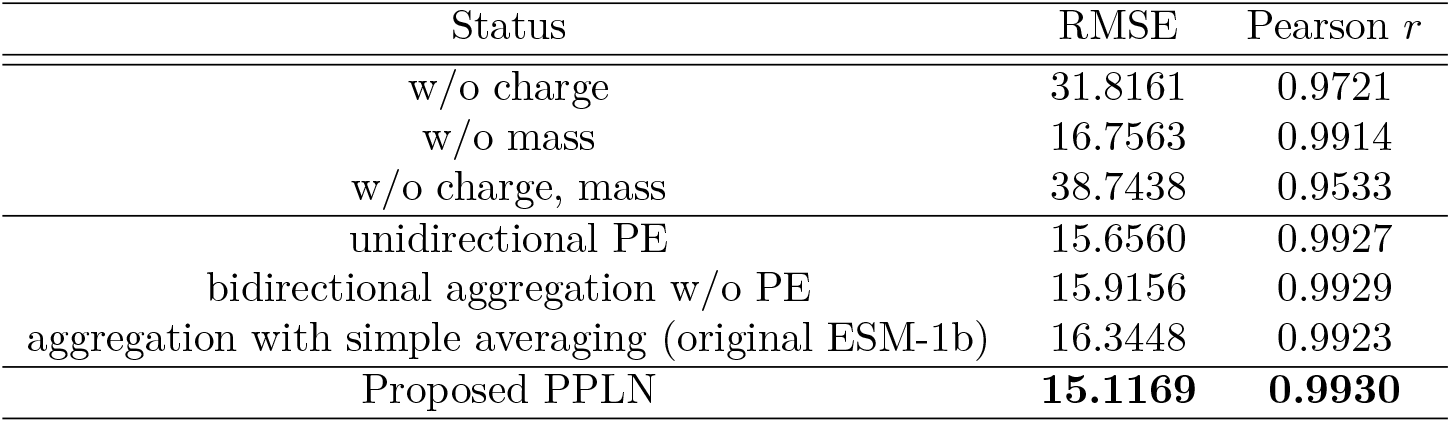
Summary of ablation study.

We first validate the necessity of including the charge number and the mass into the input of the prediction NN in the proposed PPLN. The performance of the method that does not use charge numbers of ions was much worse than the proposed method, indicating that charge number bears important information for CCS value prediction. It is in contrast with the conventional CCS prediction method [21] which does not use the charge number information. The method which excludes mass and that which excludes both charge and mass also showed worse performance. From the results, we can argue that the information of charge and mass of the ions is valuable for the CCS prediction.

We also validate the necessity of the bidirectional PE by comparing the predictive performance of the following four methods: the method with the bidirectional PE (i.e., the proposed method), the one with the unidirectional PE, the one with the bidirectional aggregation without PE (i.e., setting ***c***_*p*_ in equation (3) to be equal to the all-1 vector), and the one adopting aggregation with simple averaging (i.e., the original ESM-1b [25], setting ***c***_*p*_ in equation (4) to be equal to the all-1 vector). From Table 1, none of the compared methods showed better performance than the proposed method. The disuse of the bidirectional aggregation especially degraded the performance. This implies that the N-terminal and C-terminal sides of the ions have different effects on peptide structures and on the resulting CCS values, and those effects can be successfully learned by the network with the bidirectional aggregation.

### 2.6 Comparison of execution time for training

The proposed PPLN can simplify the training process by using the pretrained model as the feature extractor, compared with the conventional bidirectional LSTM-based method [21] that needs training from scratch. We numerically evaluated the execution time required for training of the proposed PPLN and the conventional bidirectional LSTM-based method.

Figure 6 shows time (in seconds) spent on training for each method and that required for preprocessing, i.e., feature extraction, for the proposed PPLN. We used a Linux computer with two CPUs (Intel Xeon Gold 5320, 26 cores, 2.2 GHz base clock) and 256 GB RAM. The preprocessing time means the time taken to obtain the features for all peptides in the dataset. The training time was measured when 20 %, 50 %, and 80 % of peptides in the dataset were used for training. The average time of three runs is shown in the figure. All measured values were within plus or minus 120 seconds of the shown average values. From Fig. 6, the training of the proposed PPLN not including the preprocessing was executed in 1/78, 1/30, and 1/18 of the time of the training of the conventional bidirectional LSTM-based method, when 20 %, 50 %, and 80 % samples were used for training, respectively. Even taking the preprocessing time into consideration, execution time for the proposed PPLN was reduced to 1/4 to 1/3 of that of the conventional method. Moreover, it should be noted that the proposed PPLN achieved test prediction with Pearson’s correlation coefficient over 0.99 for all runs.

**Figure 6:**
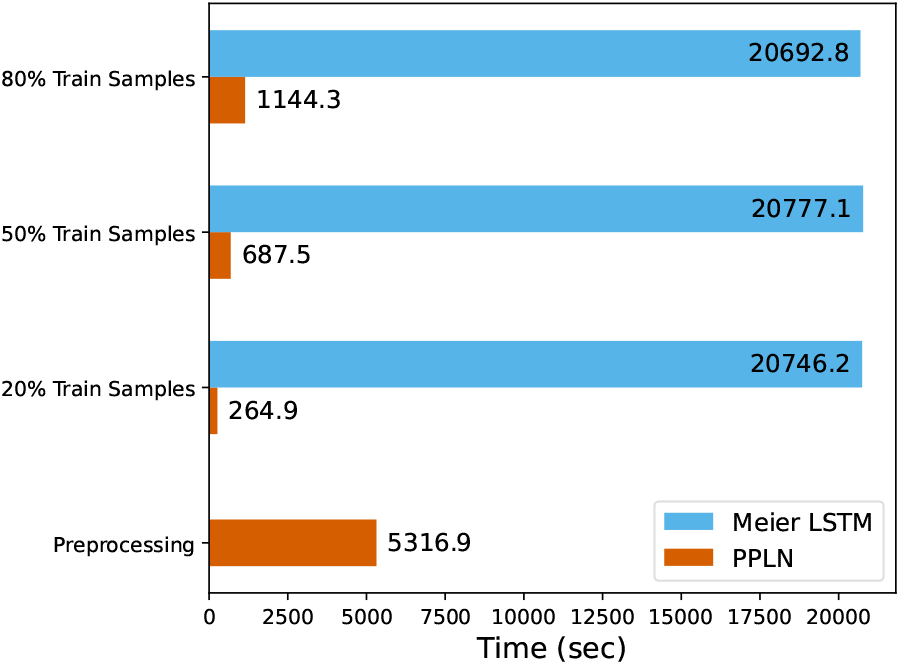
Execution time (in seconds) for the proposed PPLN and the conventional bidirectional LSTM-based method in the cases of using 20 %, 50 %, and 80 % samples for training. For the PPLN, the training time for prediction NN excluding preprocessing is shown. Execution time for preprocessing of PPLN required for obtaining features via ESM-1b is also shown separately.

These results indicate that the proposed PPLN enables a significant reduction in training time by virtue of the simplification of training through the use of the pretrained model, while maintaining high predictive performance.

### 2.7 CCS prediction for improving peptide identification

In this section, we show that accurate CCS prediction provided by the proposed PPLN allows us to improve performance of downstream tasks. We take the peptide identification task as our demonstrative example, and show that it is possible to reduce the false hits using the difference between the predicted and experimental CCS values after the candidate sequences are determined by the search engine. We used tryptic peptides from *E. coli* K12 strain BW25113 cells to verify the improvement in the identification number using PPLN, as reported previously [16]. The additional parameter, CCS error, defined as the difference between the predicted and experimental values divided by the experimental value, was used for the Mascot/Percolator approach [39]. Figure 7 shows the Venn diagram of the identification results with or without CCS error. From the figure, Percolator with CCS error identified more PSMs and stripped sequences at 1 % FDR compared with Percolator without CCS error, indicating the utility of PPLN-based CCS prediction for peptide identification in proteomics.

**Figure 7:**
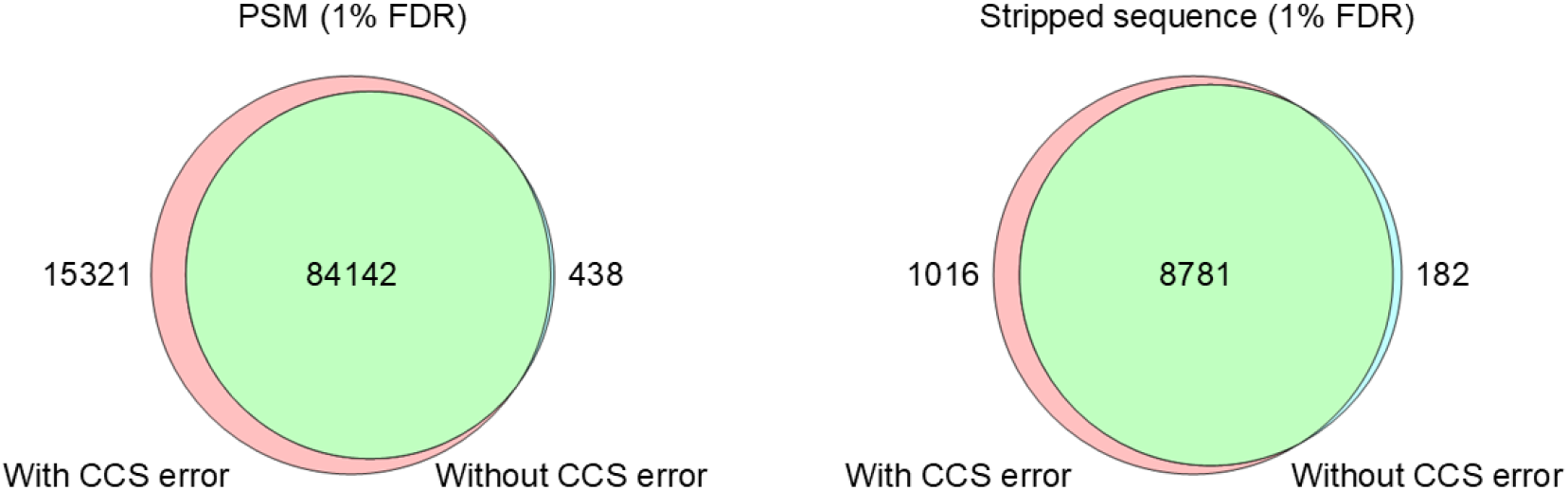
*E. coli* peptide identification results with or without CCS error.

## 3 Discussion

In this work, we proposed a novel approach to predict the CCS values of peptides using a deep learning model called the PPLN. The proposed PPLN incorporates a pretrained deep protein language model trained with a large-scale database of protein sequences as a feature extractor. Our results demonstrated that the proposed PPLN can achieve a more accurate prediction of CCS values compared with previous methods such as LS-MLR and a bidirectional LSTM-based approach. A remarkable point is that the predictions with the PPLN were made in a significantly shorter time than with the conventional bidirectional LSTM-based method requiring training from scratch. PPLN especially showed better performance in predicting CCS values of longer peptides compared with previous methods. This can be attributed to the use of the pretrained protein language model, such as ESM-1b, in PPLN: A pretrained protein language model can capture the complex relationships between amino acid sequences and protein structures more effectively than traditional methods, which might contribute to the better prediction performance of PPLN because the peptide CCS reflects the peptide structure, which is the part of the protein structure. Our study demonstrates the potential effectiveness of utilizing pretrained protein language models in predicting various peptide properties in proteomic LC/MS experiments, including not only CCS but also peptide retention time, MS/MS fragmentation pattern, and detectability. In addition to the improved prediction performance, utilizing a pretrained protein language model offers several advantages, such as requiring less training data for accurate predictions, leading to lower computational resource requirements and reducing the time and energy consumption. By utilizing these advantages, transfer learning approaches allow the model to be readily applicable to the prediction of modified peptides. It has been suggested that model size contributes to downstream task performance [40] so that the introduction of protein language models with a huge number of parameters, such as ESM-2 [27] and xTrimoPGLM [31] instead of ESM-1b used in our PPLN is expected to offer even better prediction accuracy. It would be interesting to explore the use of a larger protein language model as the feature extractor in future work.

In summary, our approach represents a significant advance in the prediction of peptide CCS values. By leveraging pretrained protein language models, we have shown that it is possible to accurately predict CCS values for longer peptides with short training times. We believe that our approach can be extended to predict other peptide properties and that the use of pretrained models will become increasingly important for efficient and accurate peptide property prediction.

## 3 Methods

### 4.1 Materials

Titanium dioxide (TiO_2_) beads were obtained from GL Sciences (Tokyo, Japan). 2-amino-2-(hydroxymethyl)1,3-propanediol hydrochloride (Tris-HCl), acetonitrile, acetic acid, ammonium bicarbonate (ABC), trifluoroacetic acid, lysyl endopeptidase (Lys-C), V8 protease (Glu-C), and other chemicals were purchased from Fujifilm Wako (Osaka, Japan). Modified trypsin and chymotrypsin were purchased from Promega (Madison, WI, USA). Asp-N was purchased from Roche diagnostics (Indianapolis, IN, USA). Lys-N was purchased from Thermo Fisher Scientific (Waltham, MA, USA). LysargiNase was purchased from Merck (Darmstadt, Germany). Alkaline phosphatase was purchased from Takata Bio Inc. (Shiga, Japan). Empore C8, Empore SDB-XC (polystyrenedivinylbenzene) and Empore SCX (strong cation exchange) extraction disks were purchased from CDS Analytical (Oxford, PA, USA). Water was purified by a Millipore Milli-Q system (Bedford, MA, USA).

### 4.2 Sample preparation

The HeLa S3 cell line (Japan Health Sciences Foundation) was cultured in 10 cm diameter dishes following standard protocols. The cells were collected, and pelleted down by centrifugation. The cell pellets were suspended in a lysis buffer, reduced and alkylated as previously reported [41]. The samples were diluted 5 times with 50 mM ABC buffer or 10 times with 10 mM CaCl_2_ in the case of LysargiNase digestion. The digestion was performed at 37*°*C by adding trypsin, Lys-C, Lys-N, Asp-N, LysargiNase, chymotrypsin or Glu-C. The appropriate enzyme-to-protein ratios were used for each enzyme (Table S1). The resulting peptides were desalted and purified via SDB-XC StageTip according to the previously published protocol [42, 43]. The desalted peptides were further processed with C8 StageTips packed with TiO_2_ to enrich phosphopeptides as previously reported [35]. Phosphopeptides were eluted with 0.5 % piperidine followed by 5 % pyrrolidine, acidified immediately by adding equal volume of 20 % phosphoric acid (final concentration: 10 %), and desalted using SDB-XC StageTips [44]. Enriched phosphopeptides were fractionated using SCX StageTips as described previously [36], followed by dephosphorylation with 6 units of alkaline phosphatase in 30 µL of 100 mM Tris-HCl buffer (pH 9.0), incubated for 3 hours at 37*°*C. After the reaction, the buffer was acidified by adding 10 % TFA 10 µL. The samples were desalted using SDB-XC StageTips.

### 4.3 LC/IMS/MS/MS analysis

LC/IMS/MS/MS was performed on an Ultimate 3000 RSLCnano (Thermo Fisher Scientific, Waltham, MA, USA) LC pump coupled with a PAL HTC-xt (CTC analytics, Zwingen, Switzerland) autosampler and a timsTOF Pro (Bruker Daltonics, Bremen, Germany) mass spectrometer. Peptides were separated on a 15 *×* cm 100 µm in-house-packed with 1.9 µm Reprosil-Pur AQ C18 beads (Dr. Maisch, Ammerbuch, Germany) column at a flow rate of 500 nL*/*min with an PRSO-V2 (Sonation, Biberach, Germany) column oven heated at 50*°*C. Mobile phases A and B were water and 20 %/80 % water/acetonitrile (v/v), respectively, both with 0.5 % acetic acid as an ion-pair reagent [45]. A total run time was 120 min with gradient starting with a linear increase from 5 % B to 40 % B over 90 min followed by linear increases to 99 % B in 1 min. The mass spectrometer was operated in data-dependent PASEF [9] mode, with 1 survey TIMS-MS followed by 10 PASEF MS/MS scans per acquisition cycle. An ion mobility scan range from 1*/K*_0_ = 0.6 to 1.5 Vs*/*cm^2^ was employed with 100 ms ion accumulation and ramp time. Precursor ions for MS/MS analysis were selected and isolated in a window of 2 *m/z* for *m/z <* 700 and 3 *m/z* for *m/z >* 700. Singly charged ions were excluded from the precursor ions according to their *m/z* and 1*/K*_0_ values. The TIMS elution voltage was calibrated linearly to obtain the reciprocal of reduced ion mobility (1*/K*_0_) using three selected ions (*m/z* = 622, 922, and 1222) of the ESI-L Tuning Mix (Agilent, Santa Clara, CA, USA)

### 4.4 Database search and data processing

MS raw files were first processed by Bruker DataAnalysis software to generate mgf files. Database search was performed with Mascot version 2.7.0 against Swissprot (downloaded on 20200318) human database containing isoforms using the appropriate digestion rules for each protease (Table S1). Carbamidomethyl (C) was set as a fixed modification, and Phospho (STY), Oxidation (M) and Acetyl (Protein N-term) were set as variable modifications. The peptide tolerance of 20 ppm and MS/MS tolerance of 0.05 Da were used. Number of 13C was set as 1 to consider monoisotopic +1 peaks as precursor ions. Percolator [39] was used for controlling the false discovery rates (FDRs) at 1 % on both the peptide spectrum match (PSM) and unique peptide level in terms of q values. The identified PSMs were further processed with IonQuant [37] (version 1.3.6) to re-assign precursor ions. The PSMs which could not be assigned to any precursor ions with IonQuant were removed for further analysis. Furthermore, the reduced ion mobility of peptide ions was calibrated linearly using three selected ions (352.33, 761.467, 933.919) which were constitutively detected in all of the raw data, to minimize the run-to-run variability of the obtained 1*/K*_0_ values. The CCS value Ω was calculated from the obtained value of 1*/K*_0_ using the Mason-Schamp equation [46]

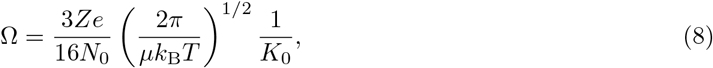

where *e* is the charge on an electron, where *Z* is the charge number of the analyte ion, where *k*_B_ is the Boltzmann constant, where *N*_0_ is Loschmidt constant, where *T* is the temperature, and where *µ* is the reduced mass of the ion and neutral given by the harmonic mean of the molecular masses of the drift gas and analyte ion, respectively. Peptide ions without any variable modifications were used for analysis. We only considered the most abundant feature for each modified peptide ion.

## 4.5 Data availability

The MS raw data and analysis files have been deposited with the ProteomeXchange Consortium (http://proteomecentral.proteomexchange.org) via the jPOST partner repository [47] (https://jpostdb.org) with the data set identifier PXD046201.

## 4.6 Code availability

The source code for CCS value prediction with the proposed PPLN is available on GitHub (https://github.com/anakai-k/PPLN).

## Acknowledgements

This work was supported by JST, CREST Grant Number JPMJCR1862, Japan. We would like to thank Genki Takahashi, a master’s student at the Graduate School of Informatics, Kyoto University, for verifying codes and formulae in this paper, and for conducting preliminary experiments.

## Author Contributions

N.-K., T. T., and Y. I. designed the experiments of the deep learning models, analyzed the data, and interpreted the results. A, N.-K. performed the experiments. Y. I. and T. T. designed the main conceptual ideas. K. O. and Y. I. collected the data. K. O. and Y. I. designed and performed the experiments of Fig. 7, and analyzed the results. All authors wrote the manuscript and discussed the results. Y. I. and T. T. supervised the project.

## S1.1 Supplementary Data

**Table S1:** Summary of digestion conditions and database search settings.

| Enzyme | E:S ratio | Missed cleavage | Cleavage rules |  |  |
| --- | --- | --- | --- | --- | --- |
|  |  |  | Sense | Cleave at | Restrict |
| Trypsin | 1:100 | 2 | C-term | KR |  |
| Glu-C | 1:25 | 5 | C-term | DE | P |
| Chymotrypsin | 1:100 | 4 | C-term | FLWY | P |
| Asp-N | 1:100 | 2 | N-term | D |  |
| Lys-C | 1:50 | 2 | C-term | K |  |
| Lys-N | 1:40 | 2 | N-term | K |  |
| LysArgiNase | 1:100 | 2 | N-term | KR | P |

**Figure S1:**
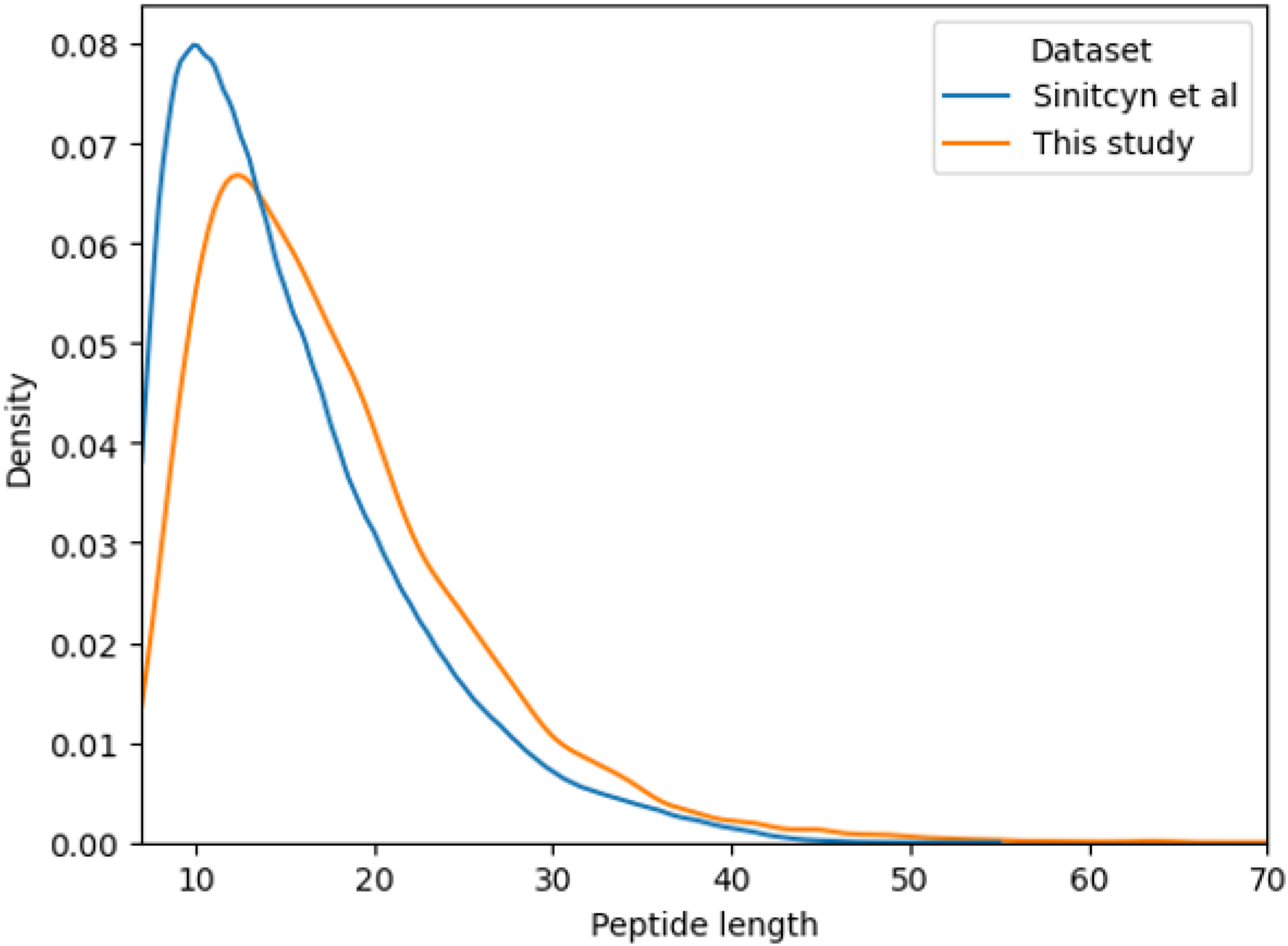
Tryptic peptide length distributions of the data sets obtained in this study and Sinitcyn *et al*. [3]. The densities were estimated using a kernel-density-based estimation function from the 31,066 and 212,798 peptide sequences for this study and Sinitcyn *et al*., respectively.

**Figure S2:**
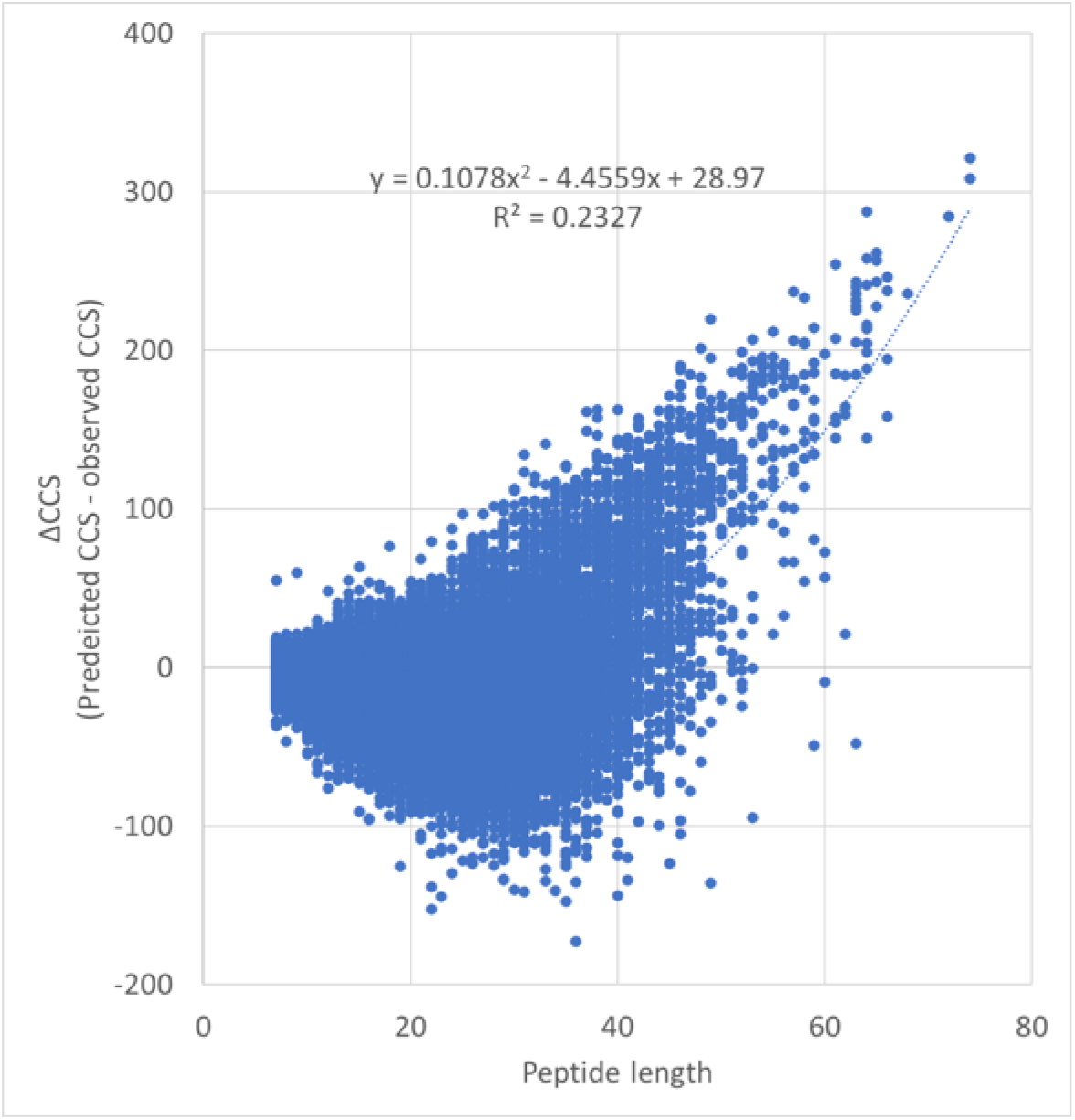
Relationship between peptide length and CCS prediction error in LS-MLR. The dashed line represents the regression line of the polynomial regression.

